# The detection of hand tremor through the characteristics of finger movement while typing

**DOI:** 10.1101/385286

**Authors:** Warwick R. Adams

## Abstract

Parkinson’s Disease (PD) is a neurodegenerative movement disease affecting over 6 million people worldwide. Current diagnosis is based on clinical and observational criteria only, resulting in a high misdiagnosis rate. Approximately 75% of people with PD have hand tremor, which can precede clinical diagnosis by up to 6 years. Previous studies have shown that early PD can be accurately detected from keystroke features while typing, and this study investigated whether tremor can be detected as well. Typing data from 76 subjects, with and without PD, including 27 with PD and 15 with tremor, was analysed and showed that hand tremor in PD can be detected from keystroke features. This novel technique has not been used before and was able to achieve an overall sensitivity of 67% and a specificity of 80% and was also able to differentiate PD tremor from essential tremor. This means that the diagnosis of early PD through typing can achieve the clinical requirement of at least two cardinal features being present (bradykinesia and tremor). Less than half a page of typing is needed, the technique does not require any specialised equipment, and can take place in the patient’s home as they type normally on a computer.

## Introduction

Parkinson’s Disease (PD) is a progressive neurodegenerative movement disease affecting 6.3 million people worldwide and is the most second most commonly occurring neurodegenerative disease in the elderly. It affects 2% of people over age 65 and the prevalence is expected to double globally over the next 20 years because of aging populations [1,2]. In PD patients, loss of dopamine-producing neurons results in a range of motor and non-motor symptoms. It has a long prodromal phase and, many years before it is diagnosed clinically, the motor signs may be detected [3,4]. However, since the symptoms of PD develop quite gradually over a 5 to 10 year period [5], the early symptoms are subtle and poorly characterised, and the current diagnostic criteria often results in a high level of misdiagnosis, not just by primary care physician (such as GP’s), but even among movement disorder specialists [6,7] and that has not significantly improved over the last 25 years. The accurate clinical diagnosis of PD depends on the presence of at least two cardinal motor features, including bradykinesia, tremor, rigidity, or postural instability [8,9].

Approximately 75% of people with PD have tremor, in particular rest tremor occurring in the frequency range of 4 – 6 Hz and being noticeable in their hand and head movement [10]. A previous study by the author [11] showed that early Parkinson’s Disease could be accurately detected using multiple characteristics of finger movement while typing. This follow-on study investigated whether hand tremor could also be detected through such finger movement and, if so, it would satisfy the diagnostic criteria of having 2 separate cardinal features present. This lead to the research hypothesis, “that hand tremors can be detected through the characteristics of finger movement while typing on a keyboard and, secondarily, that such tremors in PD patients may be differentiated from other types of tremor”.

### Tremor in Parkinson’s Disease

Tremor may be more specific than bradykinesia and it can precede clinical diagnosis of PD by up to 6 years [12]. Over the 2 years leading up to PD diagnosis, 41% of individuals report tremor symptoms to their medical practitioner compared with less than 1% of controls, and the incidence of tremor is already higher at 5 and 10 years before diagnosis [5]. Such tremor in PD occurs at a frequency of 4 – 6 Hz, prominent at the distal part of an extremity such as the hands [9], and appearing as a characteristic ‘pill rolling’ action [13]. It can also involve lips, chin, jaw and legs and, in the early stages of the disease, is usually unilateral.

Body parts generally rotate about a joint, and tremor originates in such a rotation, being detected as essentially linear motion at a distance (radius) from the joint [14]. Typing on a keyboard involves kinetic finger movement and, depending on the person’s particular typing style, they may rest their wrists on the desk surface or hold their hands unsupported above the keyboard, which may trigger postural tremor and/or dampen rest tremor. On the other hand it is recognised that, in PD patients, both attention and distraction tasks increase the power of the tremor [13], which may be due to dopamine depletion occurring during multiple tasks being unable to suppress any tremor [15]. The combination of these factors suggests that rest tremor will continue during typing.

There have been many previous studies into the detection of PD tremor using techniques based on accelerometry and gyroscopy (either with specialised sensors or smartphones) [16,17], electromyographic (EMG) signals [18], and handwriting and drawing kinematics [19]. For example, the results of a 2016 study, where users held a smartphone and executed a series of supervised FPSMT tasks [20], demonstrated accuracies greater than 85% in the rating of PD symptoms, showing promise for not just early diagnosis but also monitoring of clinical status and the effect of treatments.

Even though the kinematics of tremor with respect to wrist movement have been analysed [21] and the effect of physiological tremor on finger movement shown [22], according to literature searches, there does not appear to have been any previous investigations into the detection of such tremor through finger movement characteristics, especially during the execution of repetitive, fast movements such as typing on a keyboard.

### Significance of the research

From the perspective of patient quality of life, PD is one of the most severe of all chronic diseases and, because the most severe symptoms occur in the advanced stages of the disease, strategies aimed at early detection and treatment will have the most benefit, both for quality of life and in minimising the economic burden on health resources [23]. Because there is no definitive test for the diagnosis of PD, the initial disease diagnosis must currently be based on clinical and observational criteria only. Many symptoms of PD are imprecise and also common to other diseases, both neurodegenerative and non-neurodegenerative in nature.

Even where evaluation is performed by a physician using the Unified Parkinson’s Disease Rating Scale (UPDRS) [8,24], it is a subjective measure which leads to a lack of objectivity, sensitivity and repeatability in the scale [8,25]. Biomarkers (both biological and genetic) hold promise for reliable early PD diagnosis, while neuroimaging (SPECT) and sonography show enormous potential for high degrees of sensitivity and specificity in diagnosing early PD [7,26]. However, such diagnostic tests require a third level of medical referral – starting with the initial physician (usually a general practitioner) recognising that the presenting patient may have early PD and referring them to a neurologist, who may then get the patient to undergo specialist imaging.

The significance of this research is that it has the potential to enable provision of an objective and accurate diagnostic test for PD, based on two separate cardinal disease features (bradykinesia and hand tremors), that will overcome the need to rely entirely on the experience and skill of the physician, along with achieving an earlier diagnoses of PD, even before motor and non-motor symptoms are visually evident. It uses a novel detection technique that does not require any specialised equipment, and detection and analysis can take place as the patient types normally on their keyboard in their home environment. In addition, the technique will also be able to monitor the effectiveness of treatment (drug dosages etc.) and to measure the progression of the disease over time.

## Methods

### Data collection and processing

The tremor detection methodology was predicated on several factors - any existing hand rest tremor also being present during keyboard typing; the tremor affects the keystroke flight times between successive keys (flight time is the elapsed time between releasing one key and pressing the next); and such tremor being able to be detected from those effects, with subjects with tremor successfully differentiated from those without.

The keystroke dynamics of participants, both those with PD tremors and the controls, were captured as they typed normally on a computer keyboard throughout the day, then uploaded to a central web server database daily for analysis. The investigation included 465 anonymous participants aged 55 to 75 years (which is the age range at which the disease generally becomes evident, and hence the target range of this research), both with and without PD, and at various stages of the disease (both elapsed time since diagnosis and disease severity). The methodology utilised the same participant keystroke dataset as that for a previous investigation by the author into the detection of PD [11], but with the inclusion of keystroke data from an additional 365 participants.

The study was approved by the Human Research Ethics Committee (HREC) at Charles Sturt University, protocol number H17013, and was performed under that approval [27] and in accordance with relevant guidelines/regulations, including the informed consent of all participants and the anonymity of both participants and their data. Over the period of April to December 2017, potential research participants visited the project website, which provided details of the project, the need for volunteers and the eligibility criteria. Those interested in participating then downloaded and installed the keystroke capture application (‘Tappy’) and entered details regarding their age and disease status. For each participant, several months of keystroke data were then recorded as they typed normally on their computer (i.e. free text, rather than predetermined passages of text). The data design, keystroke capture software and overall processing flow was identical to that used in the previous study.

The characteristics of the entire participant cohort are shown in Table 1, along with the distribution of their ages and disease severity in Fig 1, however, only a subset of those was used for the subsequent analysis – those not taking levodopa medication (Sinemet^®^ and the like) in order to exclude the effect of that PD medication on their tremor characteristics; and those who had typed at least 8 sequences of 50 continuous characters or more with each hand This reduced the final sample size to 76 participants (including both PD and non-PD) and their characteristics are also listed in Table 1. It can be seen that, of the participants with PD, 56% had hand tremor, which is somewhat lower than the expected rate of 70 - 75% in the disease population. One possible reason for this could be simply that those with hand tremor do not type as much and therefore did not participate in this study.

**Table 1.** Characteristics of participant cohort.

|  |  | All Participants |  | Analysis Dataset |  |
| --- | --- | --- | --- | --- | --- |
|  |  | Count | Percent | Count | Percent |
| Disease Status | PD | 337 | 72% | 27 | 39% |
|  | Non PD | 128 | 28% | 49 | 61% |
|  | Total | 465 |  | 76 |  |
| Gender | Male | 249 | 54% | 30 | 41% |
|  | Female | 216 | 46% | 46 | 59% |
| PD Tremors | Yes | 222 | 66% | 15 | 56% |
|  | No | 115 | 34% | 12 | 44% |
| PD Severity | Mild | 196 | 58% | 18 | 67% |
|  | Medium | 116 | 34% | 6 | 22% |
|  | Severe | 25 | 8% | 3 | 11% |
| PD Sidedness | Left | 102 | 30% | 8 | 30% |
|  | Right | 151 | 45% | 14 | 52% |
|  | None | 84 | 25% | 5 | 18% |

**Fig 1.**
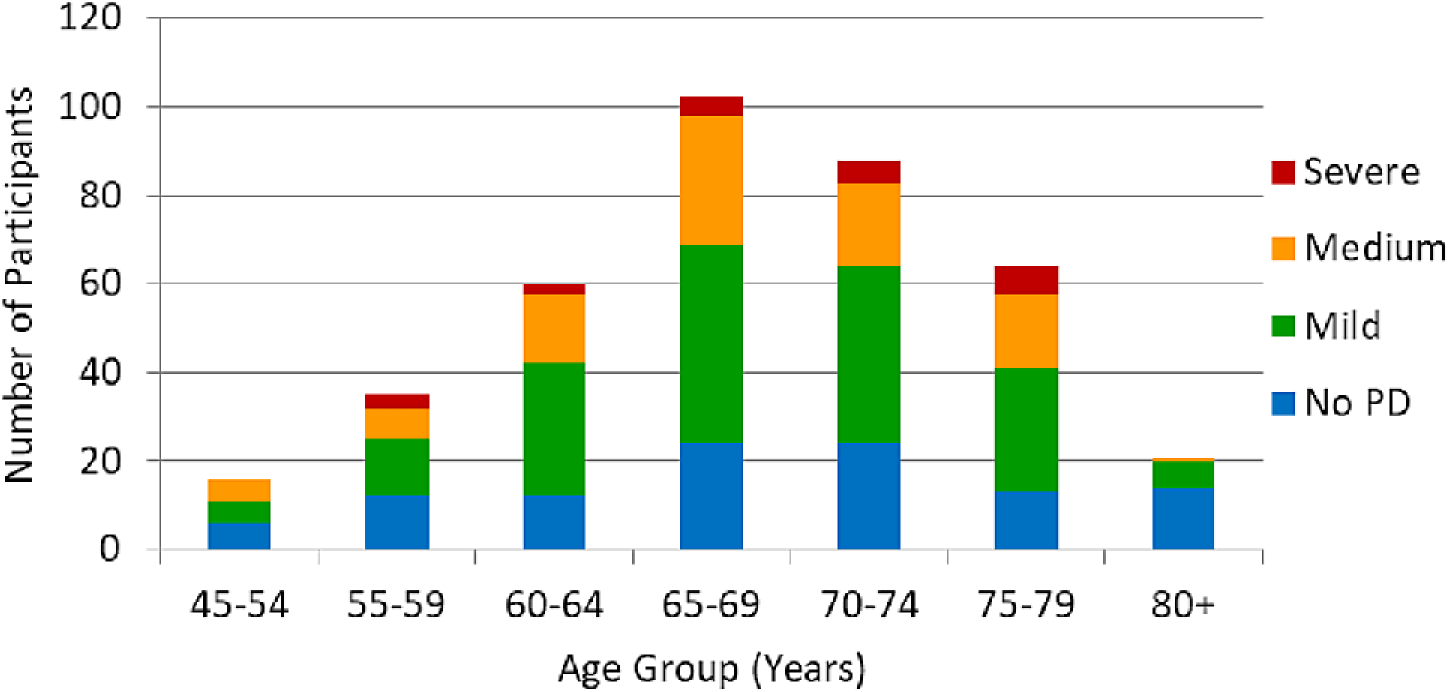
Distribution of the ages and disease status of the entire participant cohort. (all users, both taking levodopa medication & de-novo).

### Keystroke recording

The method involved monitoring all the participants’ typing, irrespective of the application they were using at the time (e.g. typing emails and documents). This design decision was made in order to facilitate data capture over a longer timeframe, as well as to avoid any stress on the participant to ‘perform well’, which could itself change their keystroke dynamics. However this decision did mean that the participants needed to install an application on their computer. The alternative of using a web-based application was decided against as, in that case, keystroke capture would be limited to just the currently-active web page, meaning that the participants would need to be led through some form of ‘typing test’ rather than being free to type normally in their various computer applications.

The Tappy application ran continuously as a background process on each participant’s PC and enable system-wide real-time recording of keystroke information. Not all keystrokes were recorded, just those corresponding to the five keyboard columns for left-hand fingers and the five for right-hand fingers, but excluding all numeric keys. The accuracy of keystroke timing data was 3.4 ± 2.0 mS (across 15 random participants with various Windows computers and keyboard types), measured according to the software delay of processing Windows keyboard events. This is consistent with the timing accuracy in another study [28] (being 3.2 ± 0.8 mS), using a similar measurement but on a known Windows computer configuration.

### Typing characteristics of participants

As already mentioned, the methodology required each participant to type continuous text segments of at least 50 keystrokes for each hand, equating to 100 keystrokes total, or approximately 15 words. Since the study design used their data as they typed normally throughout the day, the nature of that typing determined the number of such sequences available for analysis. For example, browsing the web involves just a few words, sending an email usually has complete sentences, and typing a document has larger periods of continuous typing. Fig 2 shows the actual typing patterns of the participants, both with and without PD, and it can be seen that 71% of sequences for each hand are shorter than 20 keystrokes – that is, people generally only type a short amount between pauses.

**Fig 2.**
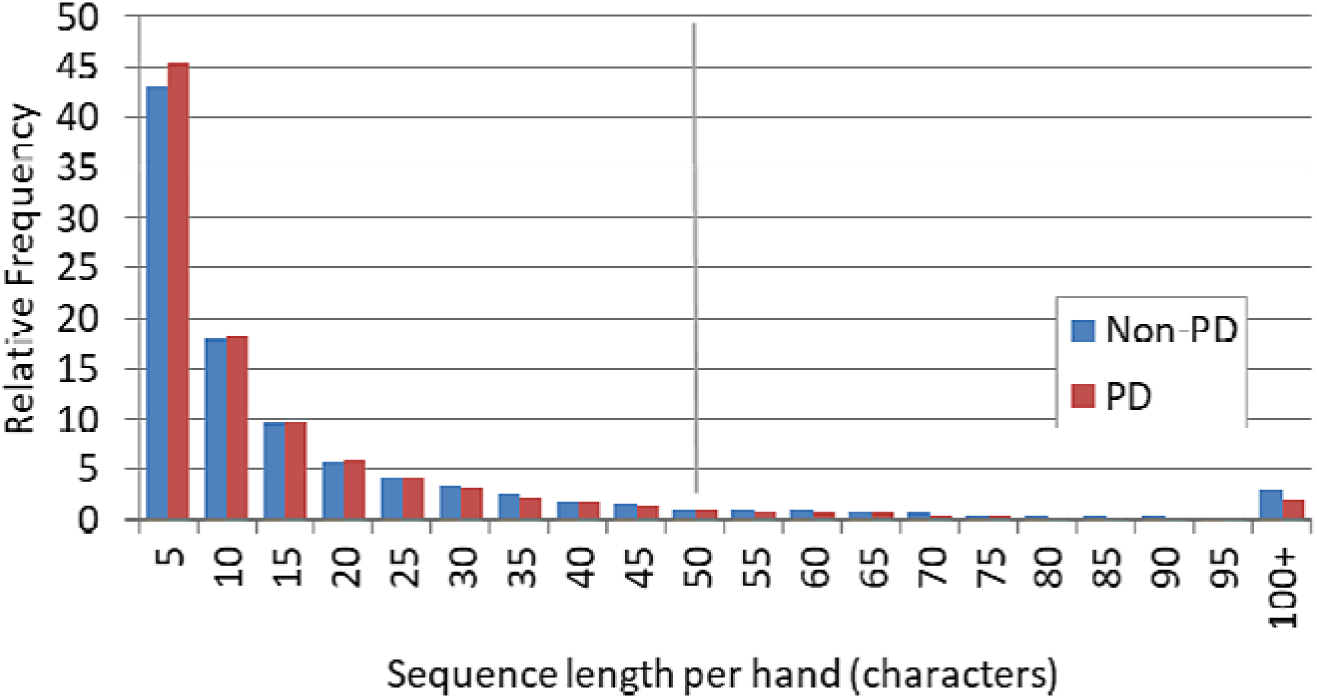
Distribution of the lengths of typing sequences. participants with PD compared to controls.

## Data analysis

A series of keystrokes and their respective flight times, after removing non-contiguous instances, forms a digital data stream with a widely spaced, non-uniform sampling interval. A Fourier analysis could, in theory, be applied to such a discrete-time signal in order to produce a frequency domain representation and to detect particular frequency components including their respective power levels indicative of hand tremors. However, in this investigation that approach was not possible because -

a. The mean sampling rate, determined by the user’s keystroke rate, was very low. For example, typing at 20 words per minute (typical for the target age range), with an average of 7.5 characters per word, is equivalent to sampling at 2.5 Hz. In addition, the data for the left and right hands needed to be separately analysed, which results in a keystroke rate for each hand of just 1.25 Hz.
b. Initial analysis of keystroke timings showed that there was an extremely broad range of keystroke rates involved.
c. As any resting tremors occur in the frequency range of 4 – 6 Hz, the minimum sampling rate required for Fourier analysis (the Nyquist frequency) would need to be around 12 Hz, an order of magnitude higher than participants’ actual keystroke rate. However, a necessary and sufficient condition for the reconstruction of a signal from irregularly spaced samples is that the density of its samples needs to exceed the Nyquist rate [29,30].
d. Investigation also found that any tremor signal component was very faint compared to the normal variations involved with pressing and releasing successive keys, with a signal to noise ratio (SNR) in the order of −40 dB to −50 dB.

This meant that quite a different technique was required in order to detect tremors from keystroke dynamics.

## Detection technique

This novel technique used a sinusoidal interpolation of the keystroke samples to test for the presence of a particular tremor frequency, by calculating the differences between each sampled flight time and the corresponding sinusoidally-interpolated value for that frequency and at the same elapsed time. The variance of all those individual differences would be at a minimum, relative to the corresponding variance at other frequencies, when a tremor signal was present at that frequency. Then, by successively applying this process to a ‘sweep’ of frequencies across the range of 3 to 10 Hz at small increments (0.05 Hz), any tremors and their frequencies could be quite accurately detected. The basic detection algorithm is shown by the pseudocode in the Appendix.

For the technique to be successful, there were also other factors to consider, such as the unknown phase angle of any tremor frequency (which was required in the analysis), the need for filtering of spurious detections (false positives) and frequency aliasing (for example, a 10 Hz signal would also appear at 5 Hz using this technique), however these could all be addressed.

### Keystroke sequences

The first step in this methodology was to extract contiguous sequences of keystrokes from the participant data, with each sequence comprising the keystrokes for just one hand (left or right). This was needed as PD hand tremors are typically asymmetrical, affecting one side more than the other (if there was a symmetrical tremor, it may have indicated Essential Tremor rather than PD). At least 50 continuous keystrokes were required in each sequence, which was terminated by either (i) 120 seconds elapsed time, or (ii) a pause of 10 seconds or more. Those constraints were required as, over a longer period of typing, or between non-contiguous sections of typing, any tremor signal was likely to be phase-shifted, masking the detection.

### Thresholds and filters

The final criteria for determining that an individual had a PD rest tremor in a particular hand was that -

a. The tremor frequency was detected across at least 4 separate replications
b. Its detection level and ‘strength’, over the 4 to 6 Hz frequency, range exceeded a predetermined threshold, calculated as –

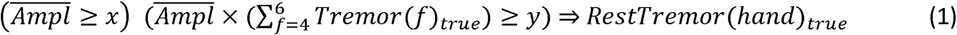

where hand is left or right, f is the frequency (Hz),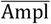 is the mean amplitude of tremor frequencies (over the 4 to 6 Hz range) for that hand and x and y are detection thresholds.

Because of the high detection sensitivity involved and the very low SNR of the signal, it was essential to apply filtering to any frequencies detected as possible tremor. Investigation of the swept frequency interpolation method showed that any possible tremor would also be detected over a small range of adjacent frequencies, typically 0.1 Hz each side of the actual frequency. This meant that, if the phase angles of the two immediately-adjacent frequencies were close to that of the detected frequency, the signal could be classified as valid, thus eliminating spurious detections.

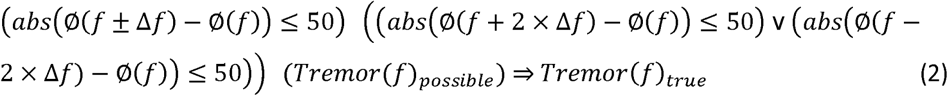

where f is the frequency (Hz) of a possible tremor signal, ∆f is a frequency offset (Hz) from f, and Ø(m) is the detected phase angle (degrees) of a signal at frequency m.

This filter proved very effective in practice, with an optimum phase angle of 50 degrees eliminating most false detections.

## Results

### Tremor detection results

A custom software application was developed to extract the features from participant’s typing data, analyse the dataset, produce tremor detection values and to display the results for individual participants in graphical form. Fig 3 shows the detection graph for a typical participant with PD tremor primarily affecting their right hand.

**Fig 3(a).**
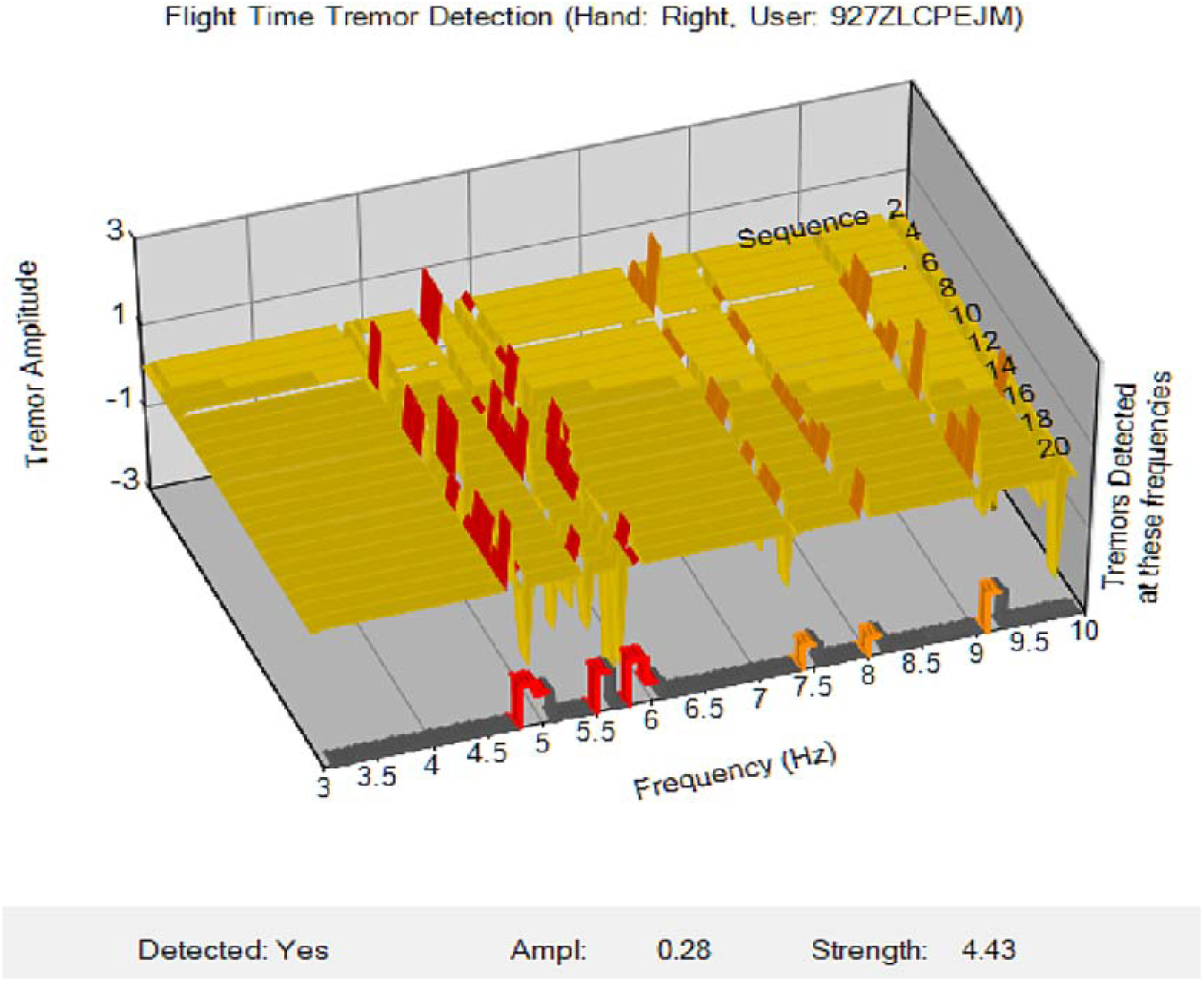
Tremor detection graph for the right hand of a participant with medium PD severity and right sidedness, showing rest tremor detected in that hand.

**Fig 3(b).**
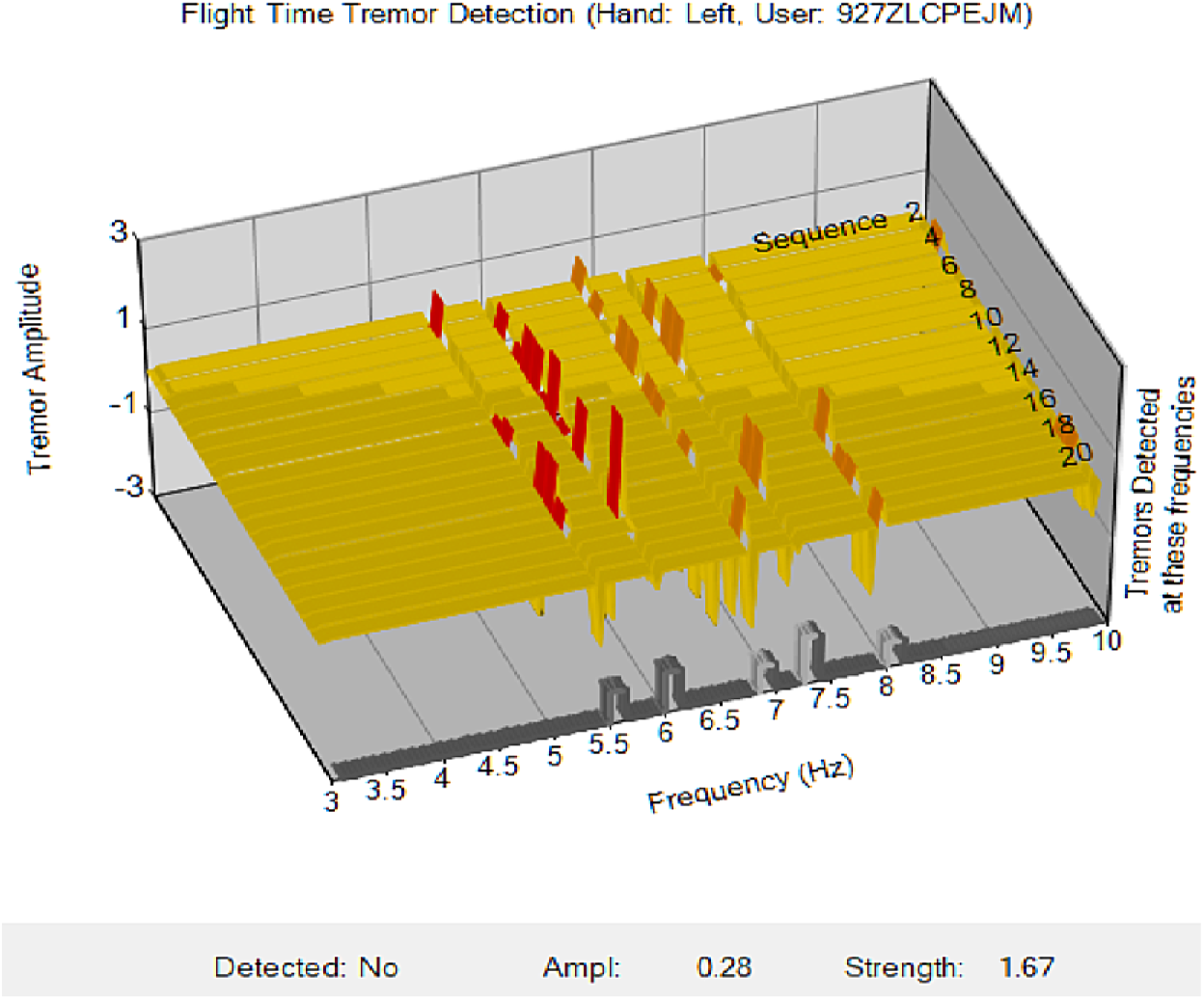
The left hand of the same participant with some indication of tremor, but below the detection threshold. The various components of the graph are as follows - X-axis, front row: Shows the frequencies of any tremors detected over the range of 3 – 10 Hz. Detected rest tremor (i.e. in the 4 – 6 Hz range) is shown in red; detected tremor frequencies outside that range are shown in orange; and where the amplitude of a possible tremor is below the established detection threshold, it is shown as grey. Y-axis: The tremor amplitude (nominal scale, related to interpolation variance). Z-axis: The individual results for 20 separate typing sequences (that is, the replications) Lower caption:Detected Yes/No. The rest tremor detection criteria were based on a combination of mean amplitude and strength thresholds. Tremor at a particular frequency must have also been detected in at least 4 of the replications. Amplitude. The mean amplitude of all rest tremor frequency detections. Strength. Number of rest tremor detections multiplied by their mean amplitude.

The tremor detection algorithm produced a 78% overall detection accuracy, with the results shown in Table 2a and the respective area under the curve (AUC) in Fig 4. This also shows that when the analysis was restricted to just include those people with mild disease severity (since the study is into detection of tremor in early PD stages), there was a notable increase in sensitivity (+6% to 73%), some increase in specificity (+2% to 82%), and the detection accuracy also improved marginally to 79%.

**Table 2.** Tremor detection results.

| a) All participants, both with & without PD |  |  |
| --- | --- | --- |
|  | All PD Severities | Mild PD Only |
| Tremor : No Tremor | 15 : 61 (total 76) | 15 : 58 (total 73) |
| PD : Non-PD | 27 : 49 | 24 : 49 |
| True positives | 10 | 10 |
| True negatives | 49 | 47 |
| False positives | 12 | 11 |
| False negatives (missed detection) | 5 | 5 |
| Sensitivity = $TP / (TP + MP) \times 100$ | 67% | 73% |
| Specificity = $(1 - (FP / (FP + TN))) \times 100$ | 80% | 82% |
| Accuracy = $((TP + TN)/Total) \times 100$ | 78% | 79% |
| b) Only participants with PD |  |  |
|  | All PD Severities | Mild PD Only |
| Tremor : No Tremor | 13 : 14 (total 27) | 9 : 9 (total 18) |
| True positives | 9 | 7 |
| True negatives | 13 | 9 |
| False positives | 1 | 0 |
| False negatives (missed detection) | 4 | 2 |
| Sensitivity | 69% | 78% |
| Specificity | 93% | 100% |
| Accuracy | 81% | 89% |

**Fig 4.**
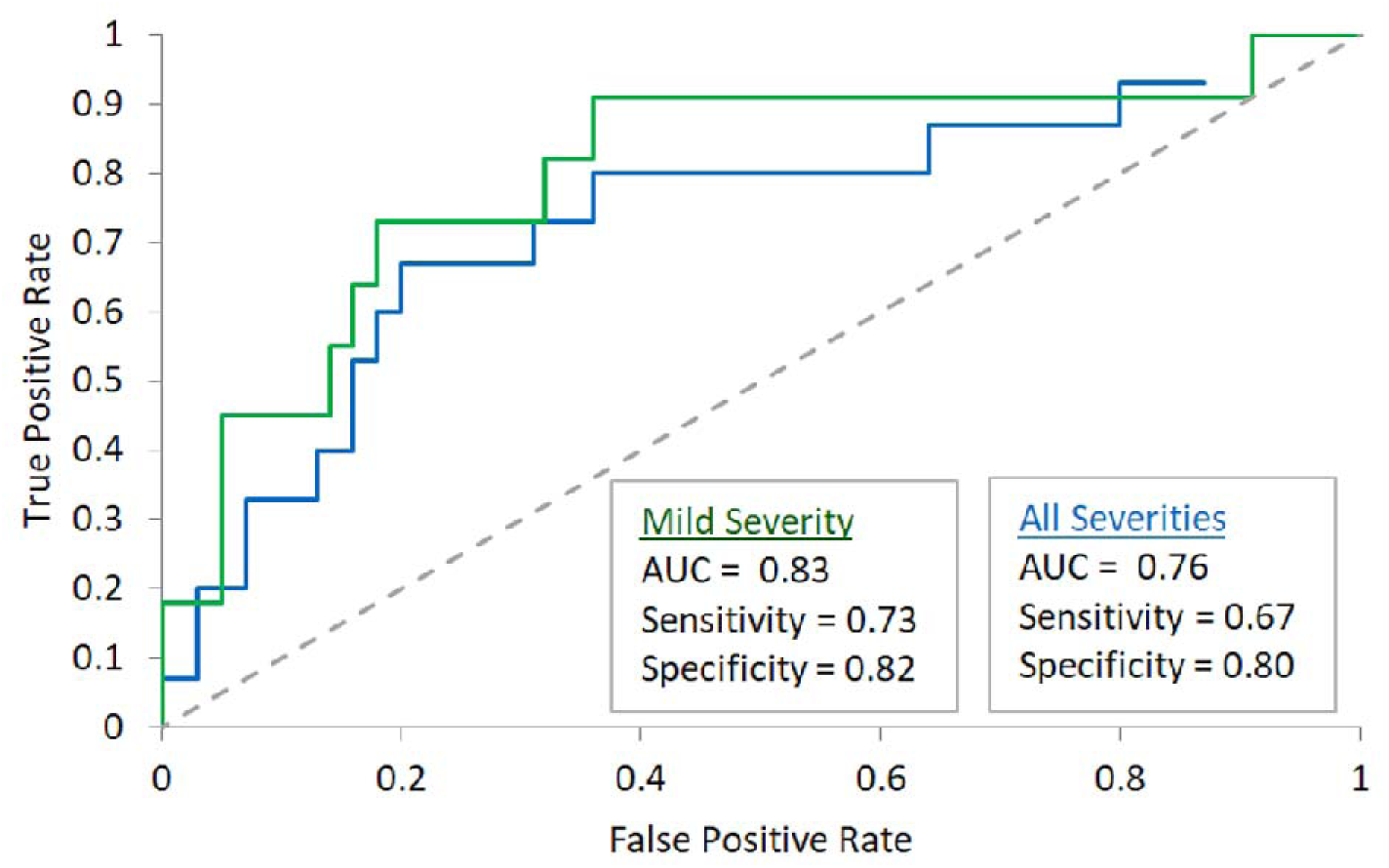
Area under the curve (AUC) results for tremor detection, controls and those with (a) any disease severity and (b) just mild severity.

The experimental design included participants both with and without tremor. There were also other variables, such as whether the participants had PD, their age, gender and possibly a range of other un-recorded conditions (arthritis for example). Since the tremor detection methodology involved analysis of each individual’s keystroke timing, and not any comparative analysis of the tremor versus non-tremor populations (such as using regression analysis or machine learning), it was considered that PD in particular would not be a confounding factor needing to be controlled in the design. Irrespective of this, it was straightforward to separate the results into ‘PD with tremor’ and ‘PD without tremor, and this was done, showing a further improvement in detection accuracy (Table 2b) - in particular, for those with mild PD severity, resulting in a sensitivity of 78%, specificity of 100% and overall accuracy of 89%.

An important criterion in this study was also to be able to differentiate between PD tremor and Essential Tremor (ET), a neurological disorder characterised by the involuntary shaking or trembling of particular parts of the body, usually the head and hands. The presence of ET is not a symptom of PD and the respective features of each are different - in particular, PD tremor (a) occurs in the frequency range of 4 – 6 Hz, rather than 5 – 12 Hz for ET, and (b) is unilateral (evident in one hand only, not both), particularly in the early stages of the disease. Using these criteria, the tremor detection technique was able to correctly discriminate between PD tremor and ET, with only one false positive in the non-PD group (comprising 49 participants).

### Data reliability and validity

With regard to this investigation, validity and reliability were assured by utilising existing participant data from a previous study, then recruiting just the additional number of participants required for the tremor analysis and for statistical validation.

The overall results were able to be verified in multiple separate ways – by cross referencing the results against the known disease, tremor, severity and medication status of the participants; by developing a confusion matrix with sensitivity, specificity and accuracy values; by statistical testing of the results, using analysis of variance (ANOVA); and forming a receiver operating characteristics (ROC) graph, where the true positive rates were plotted against false positive rates, and where the area under the curve (AUC) indicated the overall effectiveness of the method and the optimal detection cut points.

An analysis of variance, using a one-way ANOVA, showed a clear difference at the *p* < 0.05 level between the detection strength of those with tremor compared to those without tremor [*F*(1, 71) = 11.16, *p* < 0.001], as shown in Fig 5.

**Fig 5.**
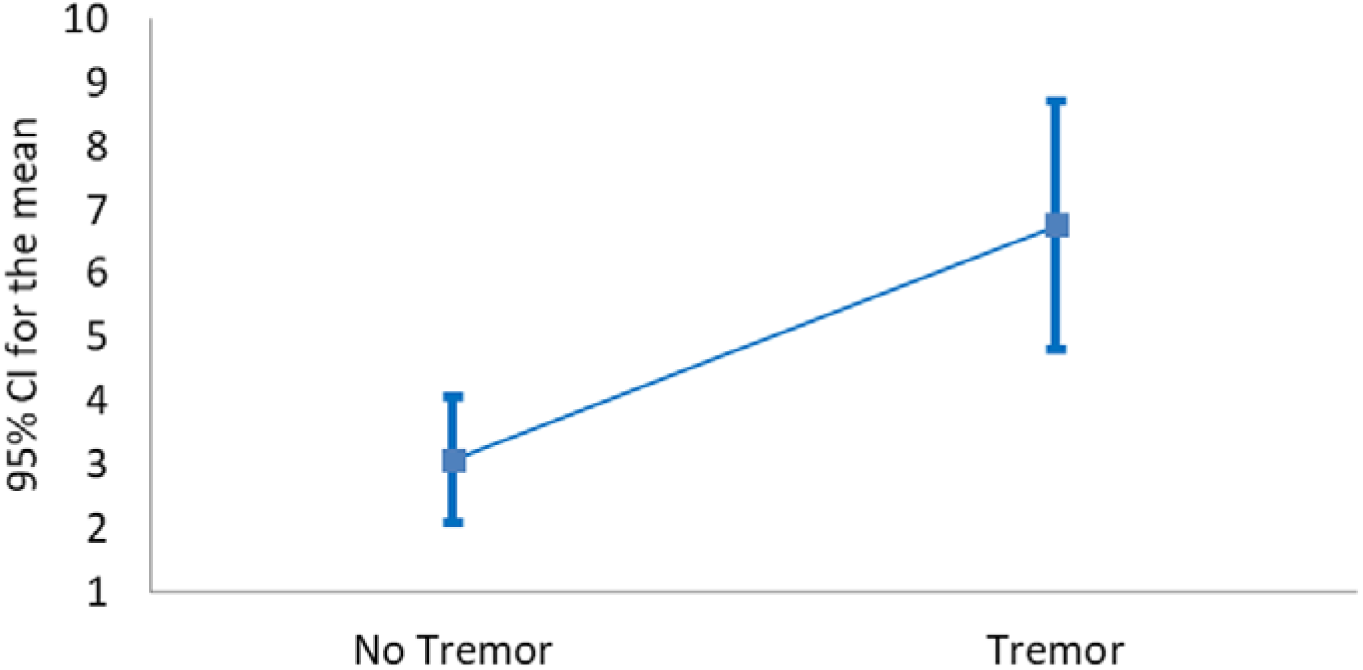
Confidence intervals for tremor detection. Participants with mild PD severity and not taking Levodopa, compared to controls at 95% confidence level.

### Effect of levodopa medication

As PD progresses in patients, most begin taking some form of levodopa medication – in essence, a form of dopamine replacement - in order to reduce the effects of the disease, and this also significantly ‘dampens’ any existing tremor (with the effect of reducing its amplitude). With respect to those study participants taking levodopa, as expected, their detected tremor strength was much lower than for those not taking it, and was similar to that of the non-tremor controls, with [*F*(1, 109) = 0.09, *p* = 0.769].

Since the purpose of the study was to detect hand tremor in people with undiagnosed PD in its early stages, those people would not be taking PD medication (levodopa prescription is generally specific to PD, not other diseases). This explains the experimental decision to exclude all participants taking levodopa from the analysis, as they would simply detract from the accuracy.

### Features and parameters

A range of thresholds were evaluated for both detection and the filtering of false positives in order to achieve maximum detection sensitivity while minimising false positives. The final criteria for determining that an individual had a rest tremor in a particular hand was (i) that the tremor frequency was detected across multiple sequences (replications), and (ii) that its detection strength, over the 4 – 6 Hz frequency range, exceeded the minimum threshold (as shown earlier in Equation 1). The parameters and cut-points established were –

> Keystroke sequences (replications) required ≥ 8
> Tremor frequency detected in at least 4 of those sequences
> Mean tremor amplitude (x) ≥ 0.15 (based on a scale relating to interpolation variance) and tremor strength (y) ≥ 4.2. These were the sensitivity thresholds.
> Phase angle filtering, requiring the phase angles of at least 3 of the 4 immediately-adjacent detection frequencies to be within 50 degrees of the tremor frequency phase angle

## Discussion

The tremor detection technique used in this investigation was successful in being able to detect PD tremor from keystroke features, without requiring any specific motor tasks to be performed by the participants. It provided a reasonable level of accuracy in the results, and was also able to discriminate between PD tremor and Essential Tremor

Although this method may not have achieved the same accuracy as some studies using wearable sensors, the objective was different and they cannot be directly compared. Wearable and attached sensors not only have limited accessibility, but also depend upon specific, supervised motor tasks being performed [31]. The keystroke data elements for this study are the same as used in a previous study by the author into detection of PD bradykinesia (the primary cardinal feature of the disease), and has shown that the same data can also be used to detect tremor, which is a second cardinal feature of PD.

There are, of course, several limitations which are inherent in the tremor detection technique used and also aspects which may provide an opportunity for future investigation, and several of these are discussed below.

### Aliasing

The nature of using sinusoidal interpolation with a swept frequency range results in aliasing, where a signal at a particular frequency will also be detected at an even division of that frequency. The technique also assumed that tremors were sinusoidal, ignoring any harmonics, which may result in a similar effect. Since the SNR of a tremor signal was so low, it was not possible to identify the primary frequency by its relative amplitude. The approach taken was to limit the valid range for rest tremor to 4–6 Hz, which means that only frequencies of 8 Hz and above would be likely to be detected as aliases. Directly filtering out possible aliases was also investigated (e.g. if a 10 Hz signal was present, to simply remove any corresponding 5 Hz signal), but that did not result in any increased accuracy.

### Limitations in detection sensitivity

In the experimental design, there were several factors which could affect the detection sensitivity, specifically the accuracy of keystroke timing, the large variance in flight times (due to the characters being typed being located in different sections of the keyboard), and the fact that a left-hand keyboard character may be pressed with a right-hand finger (and vice versa), particularly in the centremost columns of the keyboard. All of these factors simply contribute to an increased level of ‘noise’ in the data, reducing the SNR and reducing the detection sensitivity (both of a rest tremor and which side is most affected). However, the main limitation of the keystroke capture methodology used was the length of continuous sequences commonly typed by the participants. This meant that, in order to maximise the number of participants included in the analysis, the minimum sequence lengths had to be kept down to 50 characters. Analysis showed that greater detection sensitivity could be achieved with longer sequences and this could be facilitated by having participants type predetermined longer passages of text, rather than just rely on their normal patterns of typing (emails, web browsing etc.).

### Accuracy achieved and other detection techniques

This investigation has significantly progressed the definitive clinical diagnosis of PD through typing characteristics. In addition to bradykinesia, 75% of people with PD also have tremor, and 60% also have sidedness (where one side of the body is affected more than the other). All of these can be detected from keystroke characteristics, and there is potential to combine these three separate features into a single diagnostic typing test.

An alternative tremor detection technique may also be possible, by applying machine learning (ML) classification models to the features of the keystroke sequences in order to classify those with PD tremor from those without. This will be investigated further in a follow-on study, however such ML models would not identify particular frequency components, limiting their usefulness.

## Conclusion

In this investigation, keystroke timing information from 76 subjects, including 27 with PD and 15 of those with tremor, was captured as they typed on a computer keyboard over an extended period. It showed that hand tremor in PD does affect the characteristics of finger movement, which can be detected from analysis of keystroke timing features. The detection of tremor in such a manner has not been used before and a novel technique was developed, using swept frequency sinusoidal interpolation with phase angle filtering.

This technique was able to discriminate between subjects with PD tremor and those without (both non-tremor PD subjects and controls) with good accuracy, achieving a sensitivity of 67 – 73%, a specificity of 80 – 82%, and an AUC of between 0.76 and 0.83. It was also able to differentiate PD rest tremor from essential tremor with just one false positive, based on laterality and frequency characteristics.

Even though wearable sensors (accelerometers and gyroscopes) have also been shown to detect PD tremor with high accuracy, they have limited accessibility. The unique benefit of the current technique is that subjects can type normally on their own computer and the same keystroke data can also be used for detection of other cardinal symptoms of PD – bradykinesia and sidedness of movement for example.

This means that the diagnosis of early PD through the characteristics of finger movement while typing has achieved the clinical diagnostic requirement of at least two separate cardinal features being present. Less than half a page of continuous typing is needed for reliable tremor detection, the technique does not require any specialised equipment or attachments, does not need medical supervision, and can take place in the patient’s home or office environment as they type normally on a computer.

**Appendix – Pseudocode for tremor detection**

**Fig 6.**
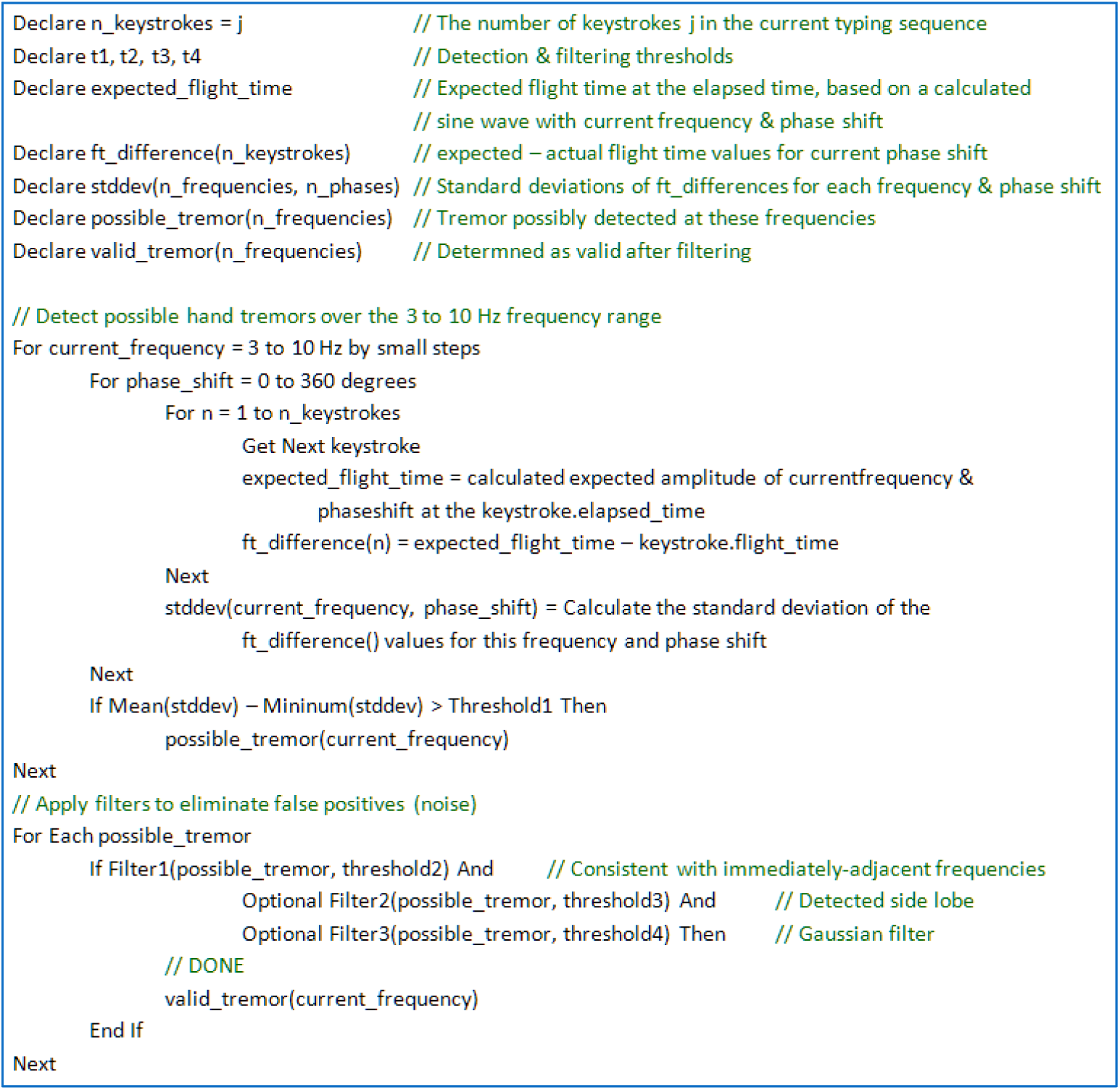
Pseudo-code for detection of a tremor, using flight times.

